# Short and long TNF-alpha exposure recapitulates canonical astrogliosis events in human induced pluripotent stem cells-derived astrocytes

**DOI:** 10.1101/722017

**Authors:** Pablo Trindade, Erick Correia Loiola, Juciano Gasparotto, Camila Tiefensee Ribeiro, Pablo Leal Cardozo, Sylvie Devalle, José Alexandre Salerno, Isis Moraes Ornelas, Pitia Flores Ledur, Fabiola Mara Ribeiro, Ana Lucia Marques Ventura, José Claudio Fonseca Moreira, Daniel Pens Gelain, Lisiane Oliveira Porciúncula, Stevens Kastrup Rehen

## Abstract

Astrogliosis comprises a variety of changes in astrocytes that occur in a context-specific manner, triggered by temporally diverse signaling events that vary with the nature and severity of brain insults. However, most mechanisms underlying astrogliosis were described using animal models, which fail to reproduce some aspects of human astroglial signaling. Here, we report an *in vitro* model to study astrogliosis using human induced pluripotent stem cells (iPSC)-derived astrocytes which replicates aspects temporally intertwined of reactive astrocytes *in vivo*. We analyzed the time course of astrogliosis by measuring nuclear translocation of NF-kB, secretion of cytokines and changes in morphological phenotypes of human iPSC-derived astrocytes exposed to TNF-α. It was observed the NF-kB nuclear translocation, increases either in the inflammation-related cytokines secretion and gene expression for IL-1β, IL-6 and TNF-α following 24 h TNF-α stimulation. After 5 days, human iPSC-derived astrocytes exposed to TNF-α exhibited increases in vimentin and GFAP immunolabeling, elongated shape and shrinkage of nuclei, which is typical phenotypes of astrogliosis. Moreover, about a 50% decrease in D-[^3^H] aspartate uptake was observed over the astrogliosis course with no evident cell damage, which suggests astrocytic dysfunction. Taken together, our results indicate that cultured human iPSC-derived astrocytes reproduce canonical events associated to astrogliosis in a time dependent fashion. Our findings may contribute to a better understanding of mechanisms governing human astrogliosis. Furthermore, the approach described here presents a potential applicability as a platform to uncover novel biomarkers and new drug targets to refrain astrogliosis associated to human brain disorders.

## 1. Introduction

Astrocytes are the most abundant cells of the mammalian central nervous system (CNS). The astrocytic processes enwrap both pre- and post-synaptic elements and closely approach the synaptic cleft, thus modulating synaptic transmission (Perez-Alvarez, Navarrete, Covelo, Martin, & Araque, 2014; Ventura & Harris, 1999). Astrocytes are also known for their secretory potential, releasing neurotransmitters, ions and other signaling molecules to the extracellular milieu in order to coordinate synaptic homeostasis (Benarroch, 2016; Verkhratsky, Matteoli, Parpura, Mothet, & Zorec, 2016).

As a cell population with heterogeneous morphology and functioning, astrocytes from rodents and humans show distinct features regarding morphology, gene expression and functional competencies (Zhang et al., 2016). Roughly, the population of human cortical astrocytes is composed of larger and morphologically more complex cells when compared to cortical astroglia from rodents (Oberheim et al., 2009). Yet, multiple subclasses of astrocytes described in the human neocortex are not found in the same region of the murine brain (Oberheim et al., 2009).

Astrocytes are prompt to react after inflammatory insults, experiencing a continuum of temporally-ordered physiological changes that culminates in a process called astrogliosis. Initially, mechanical or pathological injuries in the CNS trigger NF-kB signaling in astrocytes leading to an increased production of NF-kB-dependent cytokines, which potentially activate these cells (Lattke et al., 2017; Saggu et al., 2016). Once activated, astrocytes show progressive upregulation of intermediate filaments GFAP and Vimentin (Liu et al., 2014) and heterogenous degrees of cell hypertrophy (Kang, Lee, Han, Choi, & Song, 2014). As long as the reactive phenotype is maintained, astrocytes may present impaired metabolic functions such as disrupted recycling of neurotransmitters (Schreiner, Berlinger, Langer, Kafitz, & Rose, 2013) and energy metabolism (Gavillet, Allaman, & Magistretti, 2008). These biological events are intertwined and occur within a time course which determines the extension of the lesion (Burda & Sofroniew, 2014). However, it is unclear whether human astrocytes exhibit such response patterns since most of our knowledge about astrogliosis comes from murine systems. Currently, divergent gene expression profiles found in mouse astrocytes were activated by distinct stimuli (Hamby et al., 2012; Zamanian et al., 2012), while mouse and human astrocytes did not share the same responses to lipopolysaccharide (LPS) and IL-1 (Tarassishin, Suh, & Lee, 2014). These results highlight interspecies differences regarding astrocytic responses. Recently, murine astrocytes exposed to distinct activation stimuli were classified in A1 and A2 according to their transcriptomic profile. Interestingly, together with other cytokines, TNF-α was able to induce both profiles (Liddelow et al., 2017). Indeed, TNF-α emerges as a key player on induction, maintenance and profiling of astrogliosis.

Neural cells derived from induced pluripotent stem cells (iPSC) have increasingly been used to model human diseases. In the last few years, efforts have been made to improve differentiation protocols in order to obtain and characterize human iPSC-derived astrocytes (Chandrasekaran, Avci, Leist, Kobolak, & Dinnyes, 2016; Emdad, D’Souza, Kothari, Qadeer, & Germano, 2012; Shaltouki, Peng, Liu, Rao, & Zeng, 2013; Tcw et al., 2017; Yan et al., 2013). Owing to the uniqueness of human astrocytes as well as their responses to insults and inflammation, we evaluated human iPSC-derived astrocytes regarding the canonical events associated with astrogliosis in response to short- and long-term TNF-α stimuli as a mimetic neuroinflammatory condition. These data were obtained following a circumspect improvement of the protocol for obtaining human iPSC-derived astrocytes described by Yan and colleagues, allowing us to show that these cells present typical astroglial functional features (Yan et al., 2013). In order to provide not only a characterization of human astrocyte-secreted proteins following inflammatory stimulus, we also evaluated and characterized major events related to the time course of astrogliosis such as i) TNF-α-mediated nuclear translocation of NF-kB, ii) increases in secretion and gene expression of inflammation-related cytokines, iii) alterations in the cell shape affecting the aspect ratio and decreasing the area of astrocyte nuclei and iv) dysfunctional aspartate/glutamate uptake. Although human iPSC-derived astrocytes have been shown to respond to inflammatory stimuli (Perriot et al., 2018; Santos et al., 2017), it was unclear whether human iPSC-derived astrocytes can reproduce *in vitro* the temporally-ordered physiological changes typical of astrogliopathology.

## 2. Material and Methods

### 2.1 Generation of human iPSC-derived astrocytes

Human iPSC-derived astrocytes were differentiated from human iPSC-derived neural stem cells (NSC) obtained from iPSC of four healthy subjects. Neural stem cells (NSC) were obtained according to a protocol described previously (Yan et al., 2013). Three of these cell lines were used in other studies from our research group (Casas et al., 2018; Garcez et al., 2017). One cell line was obtained from a female subject (Subject 1 - available at Coriell Institute Biobank with the name GM23279A) and the other three from male subjects which cells were reprogrammed at the D’Or Institute for Research and Education (Subjects 2, 3 and 4). Reprogramming of human cells was approved by the ethics committee of Copa D’Or Hospital (CAAE number 60944916.5.0000.5249, approval number 1.791.182). Human cell experiments were performed according to Copa D’Or Hospital regulation. Table 1 exhibits summarized information regarding cell lines used in this study. NSC were plated at density of 5 ×10^4^ cells/cm^2^ in 75 cm^2^ culture flasks, pre-coated with Geltrex (A1413301 - ThermoFisher) in NSC expansion medium containing 50% Advanced DMEM/F12 (12634-010 - Thermo Fisher Scientific, MA, USA), 50% Neurobasal (21103-049 - Thermo Fisher Scientific, MA, USA) and neural induction supplement (a16477-01 - Thermo Fisher Scientific, MA, USA). On the following day, medium was replaced by astrocyte induction medium (AIM) composed of DMEM/F12 (11330-032 – Thermo Fisher Scientific, MA, USA), N2 supplement (17502001 – Thermo Fisher Scientific, MA, USA) and 1% fetal bovine serum (FBS) (12657029 – Thermo Fisher Scientific, MA, USA). AIM was changed every other day for 21 days. During this period, when reaching confluence, cells were passed at a ratio of 1:4 using Accutase (A6964 – Sigma Aldrich, MO, USA) to new Geltrex-coated flasks. AIM was changed on the day after cell passaging. By the end of the 21 days of differentiation, cells were exposed to astrocyte medium containing 10 % FBS in DMEM/F12. At this stage cells were named human iPSC-derived radial glia-like cells due to their positive labeling for radial glia markers PAX6 and phosphorylated Vimentin, which decrease with time in culture (Garcez et al., 2017). From then on, after reaching confluence, cells were passed at a ratio of 1:2 in the absence of Geltrex. Medium was changed twice a week and on the following day after every cell passage. Human iPSC-derived radial glia-like cells were kept in culture for additional four weeks in order to expand the number of cells and to enhance maturing of astroglial functions. All experiments described in this paper were performed from day 49 after the beginning of astrocyte differentiation (Fig. 1a).

**Table 1.** Cell lines used in the study.

| Cell Line | Gender | Age of collection | Primary Cells | Institution | Previous Publications | Use in Experiments |
| --- | --- | --- | --- | --- | --- | --- |
| Subject 1 | Female | 20 | Dermal Fibroblasts | Coriell Biobank | Casas *et al.*, 2018 | Fig. 2-6 & Supplementary Fig. 1 |
| Subject 2 | Male | 37 | Dermal Fibroblasts | D'Or Institute for Research and Education | Casas *et al.*, 2018 | Fig. 2-6 & Supplementary Fig. 1 |
| Subject 3 | Male | 31 | Dermal Fibroblasts | D'Or Institute for Research and Education | Casas *et al.*, 2018<br>Garcez *et al.*, 2017 | Fig. 2-6 & Supplementary Fig. 1 |
| Subject 4 | Female | 16 | Urine cells | D'Or Institute for Research and Education | None | Fig. 3-5 & Supplementary Fig. 1 |

**Figure 1.**
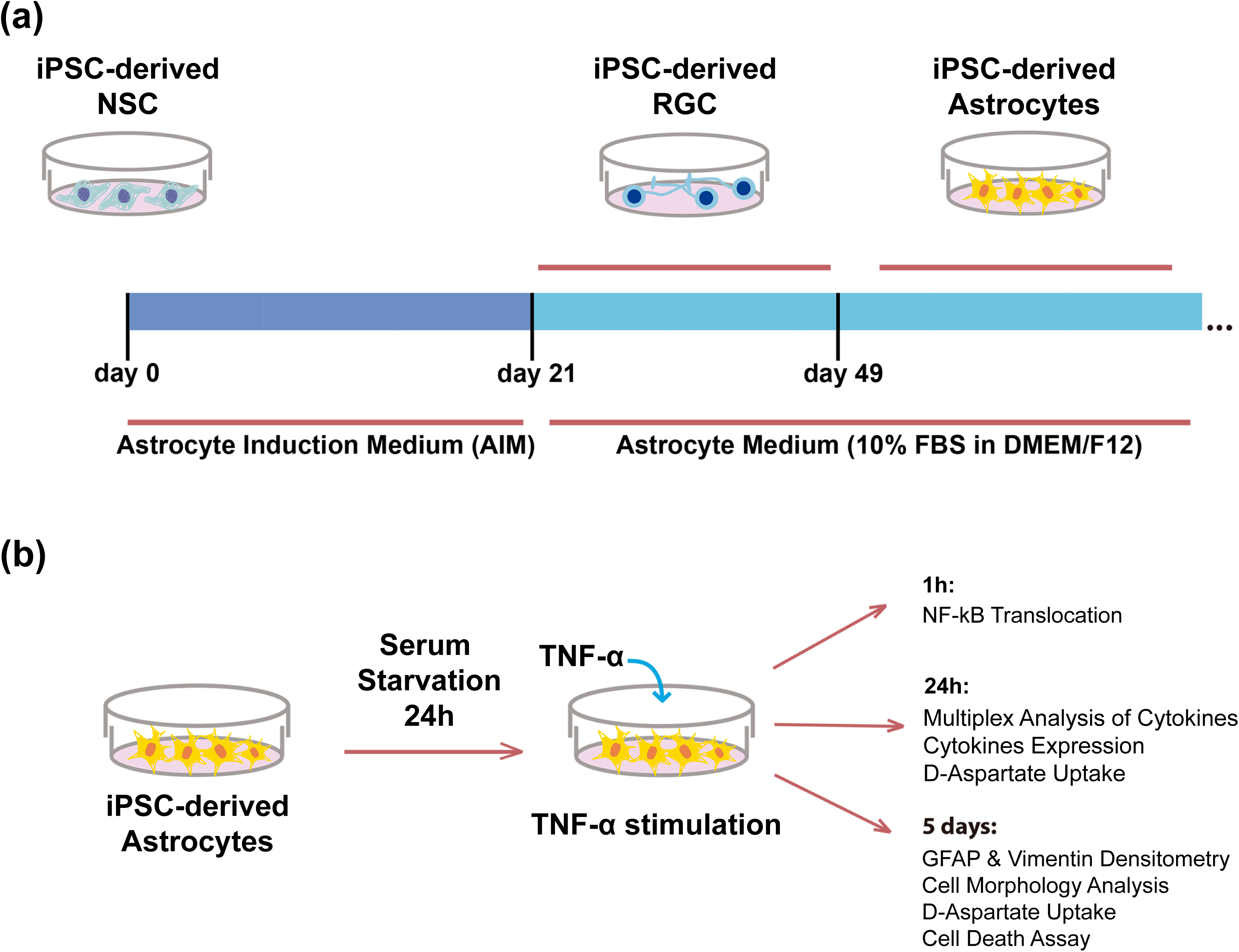
Experimental Design. **(a)** Derivation of human iPSC-astrocytes. Human neural stem cells (NSC) were differentiated from induced pluripotent stem cells (iPSC) and they were exposed to astrocyte induction medium (AIM). After 21 days in AIM human iPSC-derived radial glia-like cells (RCG) exhibited high proliferation rates and strong labeling for neural precursor markers. After exposing RGC to astrocyte medium for additional 4 weeks, human iPSC-derived astrocytes were obtained and used in the experiments. **(b)** Induction of *in vitro* astrogliosis. Human iPSC-derived astrocytes were exposed to serum-deprived culture medium for 24 h; then TNF-α was added to the cells in order to analyze the canonical events over the course of astrogliosis, as follows: i) NF-kB nuclear translocation was assessed 1 h after TNF-α exposure; ii) expression and secretion of cytokines and D-aspartate uptake assay were performed 24 h following TNF-α exposure; iii) densitometry of the intermediate filaments, vimentin and GFAP, cell morphology and viability, D-aspartate uptake assays were performed 5 days after TNF-α stimulation in order to investigate chronic stage of astrogliosis.

### 2.2 In vitro Astrogliosis

Human iPSC-derived astrocytes were seeded in non-coated multiwell plates or 75 cm^2^ flasks at densities varying from 2 × 10^4^ cells/cm^2^ to 6.25 × 10^3^ cells/cm^2^. Astrocyte medium was changed at day 1 and 3 after each passage. At day 4, astrocyte medium was replaced by serum deprived medium (DMEM/F12) for 24 h and thereafter 1 - 50 ng/ml of human recombinant TNF-α (717904 – BioLegend, CA, USA) or vehicle (DMEM/F12) were added directly to the cells. Cells were then cultured for an additional 1 h, 24 h or 5 days and samples were collected/processed/analyzed according to the experiment (Fig. 1b).

### 2.3 Immunocytochemistry and cell morphology analyses

Human iPSC-derived astrocytes were seeded at a density of 6.25 × 10^3^ cells/cm^2^ on 96-multiwell Cell-Carrier plates (PerkinElmer, MA, USA). Four days after plating, cells were exposed to serum-deprived medium and in the following day, concentrations of TNF-α ranging from 1 - 50 ng/ml were added. For NF-kB translocation experiments, cells were fixed 1 h after TNF-α exposure. For GFAP, Vimentin and morphology analyses, cells were fixed 5 days after TNF-α stimulus. Cells were fixed with 4 % paraformaldehyde (Sigma Aldrich, MO, USA) in phosphate-buffered saline for 20 minutes, permeabilized with 0.3% Triton X-100 (Sigma Aldrich, MO, USA) and then exposed to blocking solution containing 2 % bovine serum albumin (Sigma Aldrich, MO, USA). Next, an overnight incubation at 4°C with primary antibodies was performed. We have used the following antibodies in this study: mouse anti-NF-kB p65 (1:100; sc-8008 - Santa Cruz Biotechnology, TX, USA), rabbit anti-Vimentin (1:2000; ab92547 – abcam, Cambridge, UK), mouse anti-GFAP (1:200; MO15052 – Neuromics, MN, USA). Subsequently, samples were incubated with the following secondary antibodies: goat anti-rabbit Alexa Fluor 594 IgG (1:400; A-11037 - Thermo Fisher Scientific, MA, USA) and goat anti-mouse Alexa Fluor 488 IgG (1:400; A-11001 - Thermo Fisher Scientific, MA, USA) and goat anti-mouse Alexa Fluor 594 IgG (1:400; A-11032 - Thermo Fisher Scientific, MA, USA). Nuclei were stained with 0.5 µg/mL 4′-6-diamino-2-phenylindole (DAPI) for 5 minutes. Images were acquired with the Operetta high-content imaging system using 10 × and 20 × objectives (PerkinElmer, MA, USA). Total number of cells was calculated by DAPI stained nuclei counting. Densitometry and cell morphology analyses were performed using software Harmony 5.1 (PerkinElmer, MA, USA). Eleven different fields from triplicate wells per experimental condition were used for quantification.

### 2.4 Multiplex analysis of cytokines and BDNF

Cultures of human iPSC-derived astrocytes were seeded at a density of 2 × 10^4^ cells/cm^2^ in 75 cm^2^ culture flasks. Four days after plating, cells were exposed to 24 h of serum-deprived medium. Then, 10 ng/ml TNF-α was added for an additional period of 24 h. After this incubation time, conditioned medium was collected, aliquoted and immediately frozen and stored at −80°C. ProcartaPlex™ bead-based multiplex immunoassays were performed for simultaneous detection and quantitation of multiple protein targets in cell culture supernatant. The platform MAGPIX™ was used for simultaneous detection of BDNF, INF-γ, IL-1β, IL-10, IL-13, IL-2, IL-4, IL-6, IL-8 and TNF-α in a single sample according to the manufacturer’s instructions. Briefly, 50 μl/well of coated magnetic beads solution (Luminex™ Corporation) were added to a 96-well plate. The plate was inserted onto a magnetic plate washer; beads were allowed to accumulate on the bottom of each well and were then washed. Samples or standards (50 μl) were added to the wells and then incubated at room temperature for two hours on a Capp 18-X multiwell plate shaking platform (500 rpm). The plate was washed again twice and the detection antibody mixture was added and incubated at room temperature for 30 minutes on a shaking platform (500 rpm). The streptavidin solution was added to each well and incubated at room temperature for 30 minutes (500 rpm). After washing, reading buffer was added into each well and incubated at room temperature for 5 minutes (500 rpm). Data were acquired on MAGPIX™.

### 2.5 Quantitative Real time PCR

Cultures of human iPSC-derived astrocytes were seeded at a density of 2×10^4^ cells/cm^2^ in T-75 culture flasks. Four days after plating, cells were exposed to serum-deprived medium. After 24 h of serum deprivation, 10 ng/ml TNF-α was added for an additional period of 24 h. Cells were then detached with Accutase (Merck, Darmstadt, Germany), centrifuged at 300 g for 5 minutes after which the supernatant was discarded and the resulting pellet was submitted to RNA extraction. Total RNA was isolated using TRIzol™ reagent, according to the manufacturer’s instructions (Thermo Fisher Scientific, MA, USA). The total RNA was resuspended in a final volume of 12 µl of Nuclease-free water and quantified by absorbance at 260 nm in a spectrophotometer (NanoDrop 2000, Thermo Fisher Scientific, MA, USA). One µg of total RNA was reverse-transcribed in a 20 µl reaction volume using M-MLV (Thermo Fisher Scientific, MA, USA), the generated cDNA was diluted 5x and a quantitative PCR was performed using Eva™ Green PCR Mix (Biotium, CA, USA) in the StepOne Plus Real-Time PCR Platform (Applied Biosystems, CA, USA). RT-qPCR was carried out to detect transcripts of the following genes: tumor necrosis factor α (TNF-α; forward: 5’-CTGCACTTTGGAGTGATCGG-3’; reverse: 5’-TGAGGGTTTGCTACAACATGGG-3’); interleukin 1β (IL-1β; forward: 5’-CACGATGCACCTGTACGATCA-3’; reverse: 5’-GTTGCTCCATATCCTGTCCCT-3’); interleukin 6 (IL-6; forward: 5’-TACCCCCAGGAGAAGATTCC-3’; reverse: 5’-GCCATCTTTGGAAGGTTCAG-3’); glyceraldehyde-3-phosphate dehydrogenase (GAPDH; forward: 5’-GCCCTCAACGACCACTTTG-3’; reverse: 5’-CCACCACCCTGTTGCTGTAG-3’); hypoxanthine phosphoribosyltransferase 1 (HPRT-1; forward 5’-CGTCGTGATTAGTGATGATGAACC-3’; reverse: 5’-AGAGGGCTACAATGTGATGGC-3’); ribosomal protein lateral stalk subunit P0 (RPLP0; forward: 5’-TTAAACCCTGCGTGGCAATC-3’; reverse: 5’-ATCTGCTTGGAGCCCACATT-3’); and importin-8 (IPO8; forward: 5’-TCCGAACTATTATCGACAGGACC-3’; reverse: 5’-GTTCAAAGAGCCGAGCTACAA-3’). All primers used in this study were validated by serial dilution to allow reaction efficiency calculation. Efficiencies from all tested primers ranged from 90%-110%, and the maximum allowed difference in efficiencies for primers used to estimate fold change of each gene was 10% (data not show). RT-qPCR data was calculated by the 2-ΔCt method using the geometric mean of RPLP0, GAPDH and IPO8 (IL-1β) or GAPDH and HPRT1 (IL-6) to normalize results.

### 2.6 D-Aspartate uptake

Cultures of human iPSC-derived astrocytes were seeded at a density of 2×10^4^ cells/cm^2^ on 24-well plates. Four days after plating, cells were exposed to serum-deprived medium. After 24 h of serum deprivation, 10 ng/ml TNF-α was added for an additional period of 24 h. Cultures were then incubated with 1 μCi/ml of 2,3-[^3^H]-D-aspartate (11.3 Ci/mmol) in Hanks saline solution (HBSS) for 15, 30 and 60 minutes. The competitive inhibitor of the excitatory amino acid transporters (EAATs) family, 100 μM DL-threo-β-Benzyloxyaspartic acid (DL-TBOA), was added 10 minutes prior to 2,3-[^3^H]-D-aspartate. After incubating periods, uptake was terminated by washing cells with HBSS in order to remove non-incorporated 2,3-[^3^H]-D-aspartate, followed by cell lysis. Aliquots were taken for intracellular radioactivity quantified by scintillation counting. Experiments were performed with 2 replicates per experimental group in all time points analyzed and the average value of both replicates was used for statistical analyses.

### 2.7 Cell death assay

Cell death was assessed 5 days after exposing cells to 10 ng/mL TNF-α exposure. Cells were loaded with the fluorescent dye ethidium homodimer (2 µM-LIVE/DEAD Cell Imaging Kit, Thermo-Fisher, MA, USA) for 30 min. Live cell images were acquired with the Operetta high-content imaging system using a 20x objective (PerkinElmer, MA, USA). Cell death was determined by the ratio of ethidium homodimer labeling per DAPI labeling. High-content image analysis Harmony 5.1 was the software used for data analysis (PerkinElmer, MA, USA). Seven different fields from duplicate wells per experimental condition were used for quantification.

### 2.8 Statistical Analyses

Statistical comparisons between two experimental groups were performed by Unpaired Student t-test. NF-kB translocation experiments were analyzed by One-way ANOVA followed by Tukey’s post hoc test. Aspartate uptake experiments were analyzed by One-way ANOVA followed by Dunnett’s post hoc test. Statistical significance was considered for P < 0.05. GraphPad Prism v8.02 (GraphPad Software, CA, USA) was the software used for data analyses and graphics.

## 3. Results

### 3.1 TNF-α induces NF-kB nuclear translocation in iPSC-derived astrocytes

First, we aimed to determine whether human iPSC-derived astrocytes were responsive to TNF-α by analyzing nuclear translocation of the transcription regulator NF-kB, a mandatory step on astrogliosis induction. It can be noted that cells incubated for 1 hour with TNF-α ranging from 1 to 50 ng/mL showed nuclear translocation of NF-kB in all tested conditions (Fig. 2a). The increase of NF-kB in the nucleus was concentration-dependent, with 1, 5, 10, 20 and 50 ng/ml TNF-α promoting increases of NF-kB immunoreactivity around 97%, 138%, 156%, 186% and 184%, respectively (Fig. 2b). When the whole cell area was analyzed, a trend toward increasing was observed for NF-kB in all concentrations of TNF-α tested, although it did not reach statistical significance (Fig. 2c). The translocation index of NF-kB (calculated by immunoreactivity of nuclei/cell area ratio), revealed that TNF-α induced a significant increase in the NF-kB translocation index in all concentrations tested. Similar to nuclei NF-kB immunoreactivity, the increase was dose dependent, as 1, 5, 10, 20 and 50 ng/ml of TNF-α promoted an increase of NF-kB nuclear translocation around 74%, 110%, 106%, 130% and 138%, respectively (Fig. 2d). Despite the fact that cells were responsive to 1 ng/ml TNF-α for NF-kB nuclei translocation, cells that received 10, 20 or 50 ng/ml TNF-α were even more effective on recruiting NF-kB to the nucleus. Based on these findings, 10 ng/mL TNF-a was chosen for further experiments in order to promote astrocytes activation.

**Figure 2.**
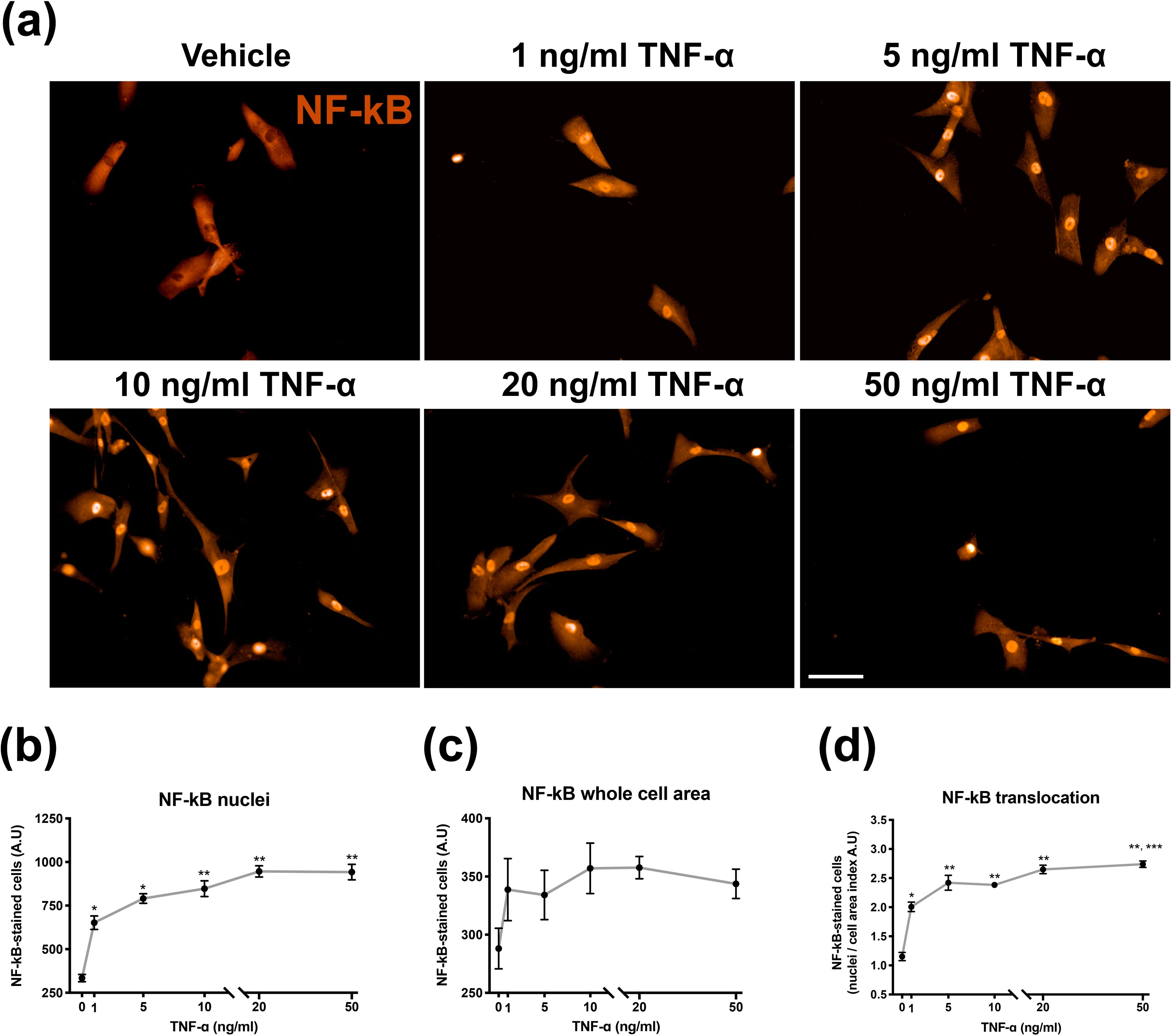
TNF-α-induced NF-kB nuclear translocation in human iPSC-derived astrocytes. **(a)** - Photomicrographs of NF-kB immunostaining 1 h after exposing cells to vehicle or five different concentrations of TNF-α. Quantification of NF-kB immunoreactivity in both **(b) –** Nuclei and **(c)** – Whole cell area, which were expressed in arbitrary units of immunofluorescence (A.U.). **(d)** – The NF-kB translocation index (nuclei/whole cell area ratio). Data are presented as means ± SEM of experiments performed in triplicates from 3 cell lines. *P < 0.01, different from vehicle; **P < 0.01, different from vehicle and 1 ng/mL TNF-α *** P < 0.01, different from 10 ng/mL TNF-α. One-way ANOVA followed Tukey’s post hoc test. Magnification: 100 ×. Calibration bar: 200 μm.

### 3.2 Inflammation-related cytokines are increased after TNF-α stimulation

The release of cytokines is a hallmark of inflammatory response and also astrogliosis. Therefore, in order to map the secretion of immunomodulators by human iPSC-derived astrocytes activated by TNF-α, we performed a multiplex analysis of cytokines using conditioned media from these cells. Based on the literature, cytokines were classified into three major categories related to their primary biological roles on inflammation: pro-inflammatory, modulatory or anti-inflammatory, and also brain-derived neurotrophic factor (BDNF). The secretion of pro-inflammatory cytokines IL-1β, IL-8 and IFN-γ was significantly stimulated by TNF-α. While IL-1β secretion was increased in 425% (from 12.32 ± 1.11 to 64.72 ± 14.05 pg/mL) and 254% for IL-8 (from 1.09 ± 0.16 to 3.85 ± 0.39 ng/mL), IFN-γ presented 890% increase (from 47.9 ± 22.5 to 474.3 ± 147.8 pg/ml) when compared to conditioned media from vehicle-treated cells (Fig. 3a). As expected, IL-1β, IFN-γ and TNF-α were barely detectable in conditioned media of non-stimulated human iPSC-derived astrocytes. TNF-α was highly detected in the conditioned media from stimulated cells, probably due to its remaining content from the activation stimulus.

**Figure 3.**
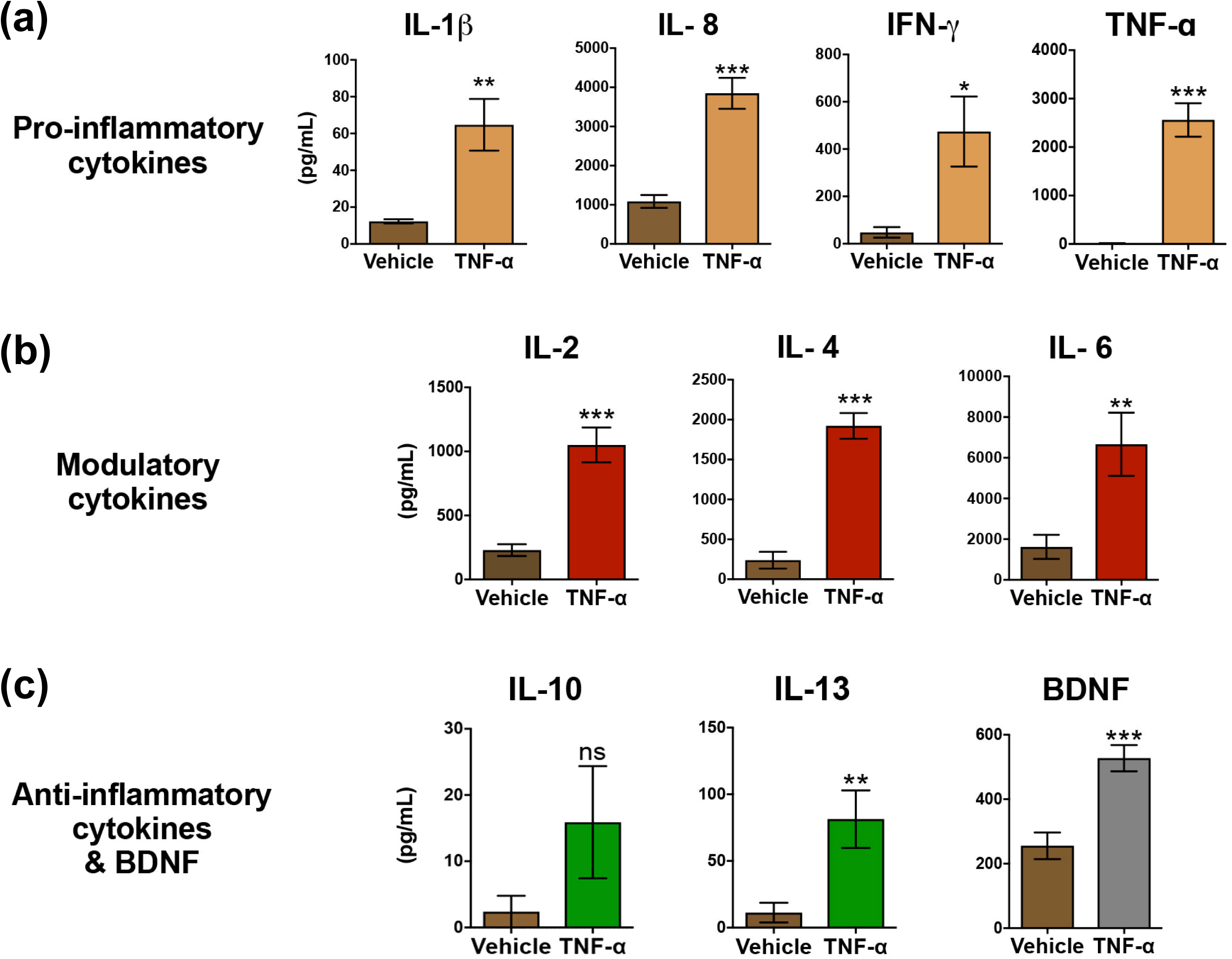
Cytokines and BDNF secretion from human iPSC-derived astrocytes. Conditioned media were collected after stimulating cells during 24 h with 10 ng/mL TNF-α. Cytokines and BDNF secretion was measured in the conditioned media and compared with cells treated with vehicle. **(a)** - Pro-inflammatory cytokines: Interleukin-1 beta (IL-1β), Interleukin-8 (IL-8), Interferon gamma (IFN-γ) and Tumor necrosis factor alpha (TNF-α); **(b)** - modulatory cytokines: Interleukin-2 (IL-2), Interleukin-4 (IL-4) and Interleukin-6 (IL-6); **(c) -** anti-inflammatory cytokines Interleukin-10 (IL-10), Interleukin-13 (IL-13) and Brain-derived neurotrophic factor (BDNF). Data are presented as means ± SEM of concentrations in (pg/ml) of secreted factors. Conditioned media were collected from 4 cell lines and the experiments were performed in duplicates. *P < 0.05; **P < 0.01; ***P < 0.001; ****P < 0.0001; ns - non-significant. Unpaired Student’s t-test.

Regarding modulatory cytokines, conditioned media obtained after TNF-α stimulus presented a 352% increase for IL-2 (from 0.23 ± 0.05 to 1.05 ± 0.14 ng/mL), 704% for IL-4 (from 0.24 ± 0.11 to 1.92 ± 0.16 pg/mL) and 311% for IL-6 secretion (from 1.62 ± 0.59 to 6.67 ± 1.55 pg/ml) (Fig. 3b). Among anti-inflammatory cytokines, while IL-10 levels were unchanged, secretion of IL-13 increased 618% (from 11.3 ± 7.4 to 81.4 ± 21.5 pg/mL) and BDNF increased 106% in the conditioned medium after TNF-α exposure (from 255.5 ± 41.4 to 527.5 ± 40.6 pg/mL) (Fig. 3c).

### 3.3 Inflammation-related cytokines display increased expression after TNF-α stimulation

Despite the fact that most cytokines have short half-life, increases in their secretion should impact gene expression and continuous production was expected. Thus, quantitative real time PCR (qPCR) was conducted to verify the expression of cytokines, which secretion was found to be increased in the conditioned medium (Fig. 4). Indeed, both IL-6 and IL-1β genes expression showed ∼15.6- and ∼ 6.0-fold increase, respectively, following TNF-α exposure (Fig. 4a,b). In addition, TNF-α expression was detected only in stimulated cells, but not in any replicates within the control group (Fig. 4c). In order to show that control samples were viable in terms of gene expression, we also analyzed the expression levels of housekeeping genes RPLP0, GAPDH and IPO8 from the same samples used in Figures 4b and 4c. No differences were found among experimental groups (Supplementary Figure 1).

**Figure 4.**
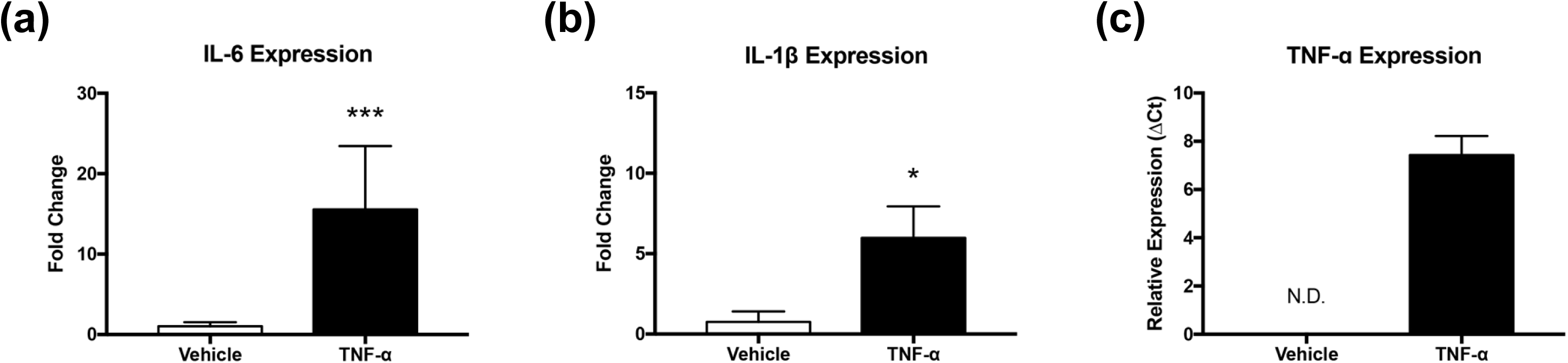
Cytokines expression in cell extracts from human iPSC-derived astrocytes. **(a)** – Interleukin-6; **(b)** – Interleukin-1 beta and **(c)** – TNF-α expression was assessed 24 h after exposing cells to vehicle or 10 ng/mL TNF-α. Data are presented as means ± SEM of fold change for (a) and (b) and ΔCt (c) since the basal levels of TNF-α were below the limit of quantification in non-stimulated cells. Experiments were performed in triplicates from 4 cell lines. *P < 0.05; ****P < 0.0001; N.D. non detected. Unpaired Student’s t-test.

### 3.4 TNF-α induced morphological alterations and upregulation of intermediate filaments Vimentin and GFAP

The incubation of astrocytes with 10 ng/ml TNF-α during five consecutive days (in serum-free conditions) triggered typical phenotypes of astrogliosis, as revealed by increased immunoreactivity for Vimentin and GFAP (Fig. 5a, b, c) as well as shrunken nuclei (Fig. 5f). The incubation with 10 ng/ml TNF-α did not change measurements of cell area (Fig. 5c). However, some vimentin-stained astrocytes clearly showed a thin, process-devoid and polarized phenotype after treatment with 10 ng/ml TNF-α. To identify these cells, we calculated the aspect ratio (length to width ratio) from vimentin immunoreactivity. A greater length/width ratio would indicate cells with relatively thin shapes (Fig. 5f). We confirmed that TNF-α promotes an increase in cell polarization (25%), which is typically found under inflammatory conditions (Jang et al., 2013).

**Figure 5.**
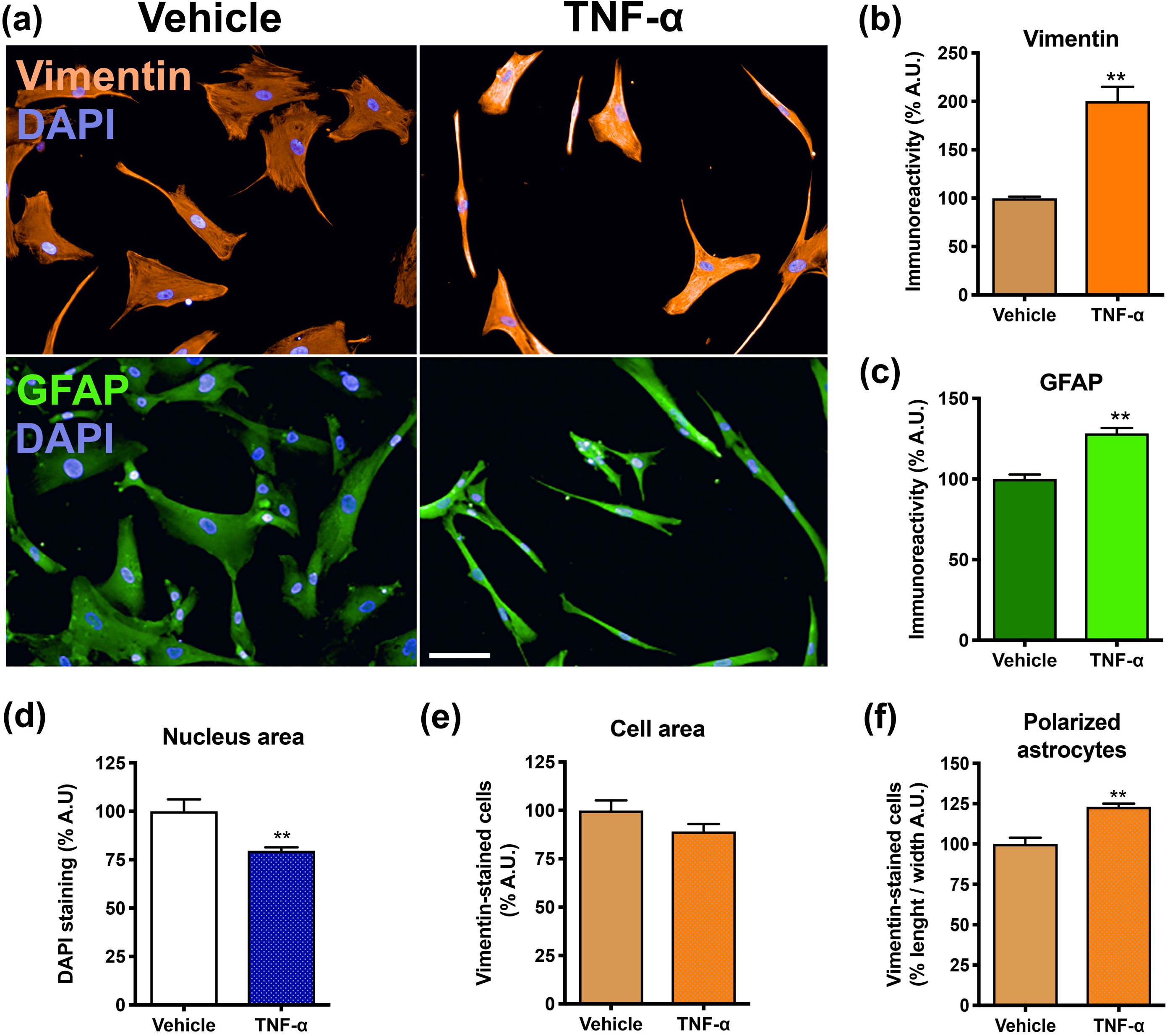
Morphological analysis of human iPSC-derived activated astrocytes following 5 days TNF-α stimulation. **(a)** - Photomicrographs of human iPSC-derived astrocytes immunostained for vimentin (red), GFAP (green) and DAPI (blue); **(b)** – Quantification of vimentin immunostaining; **(c)** – Quantification of cell area in vimentin-stained cells. **(d)** – Percentage of astrocyte polarization, which cells were classified according to increase of length/width ratio of vimentin-stained labeling. **(e)** - Quantification of GFAP immunostaining expressed in arbitrary units of immunofluorescence (A.U); **(f)** - Quantification of DAPI-stained nuclei areas expressed as percentage of micrometers. Data are presented as means ± SEM from 4 cell lines in experiments performed in triplicates. **P < 0.01. Unpaired Student’s t-test. Photomicrographs magnification: 200x. Calibration bar: 100 μm.

### 3.5 Glutamate uptake is impaired in human iPSC-derived astrocytes exposed to TNF-α

One important functional feature of astrogliosis is the dysfunction of glutamate/aspartate transporters. In order to accurately measure uptake activity, the non–metabolized analog D-[^3^H] Aspartate was used. Aspartate uptake was carried out 1 and 5 days after TNF-α incubation. At both time points, TNF-α was able to inhibit D-[^3^H]aspartate uptake by human iPSC-derived astrocytes (Fig. 6a,b). At 60 minutes of aspartate uptake, a reduction of ∼47% was found in cells exposed to TNF-α for 1 day (Fig. 6a). When cells were exposed to TNF-α for 5 days, a reduction in aspartate uptake was observed at 5, 15, 30- and 60-minutes time points (61.5%, 72.4%, 73.3% and 51.5%, respectively) when compared to control cells (Fig. 6b). Also, human iPSC-derived astrocytes displayed a highly specific aspartate uptake activity, as evidenced by full inhibition with the competitive inhibitor of excitatory amino acid transporters DL-TBOA (Fig. 6a, b). In the presence of DL-TBOA, TNF-α had no effect on transporters activity (Fig. 6a, b). Since the astrogliosis induction was carried out in the absence of serum, we assessed astrocyte viability after five consecutive days of TNF-α exposure. The removal of fetal bovine serum from culture medium induced a small increment in cell death: 5% of cells incorporated ethidium bromide while only 0.7% were ethidium-positive among serum-supplemented cells (Fig. 6c). Similarly, TNF-α in the absence of serum decreased astrocyte viability in 7.4%. No significant difference in cell death was observed between cells exposed to TNF-α and vehicle (Fig. 6c).

**Figure 6.**
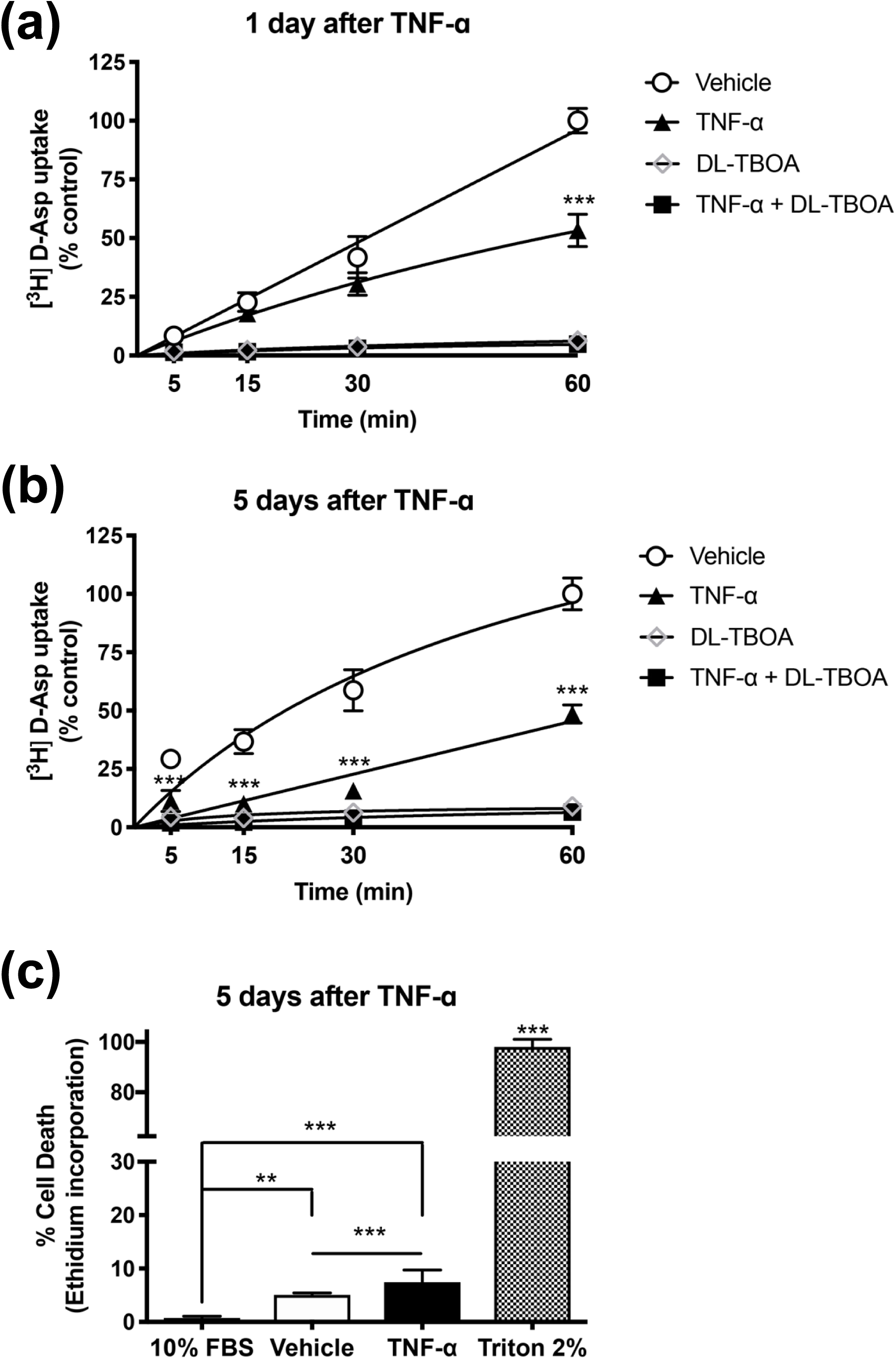
Impairment of [^3^H] D-aspartate uptake by TNF-α in human iPSC-derived astrocytes. Aspartate uptake was carried out **(a)** – 1 day or **(b)** – 5 days after exposing cells to vehicle or 10 ng/mL TNF-α. The competitive inhibitor of glutamate transporters DL-TBOA was added 10 minutes prior to aspartate. Data are presented as means ± SEM of the percentage of counts per minute (cpm). **(c)** – Cell viability was evaluated by ethidium incorporation. As a positive control for cell death, cells were lysed with Triton 2 %. As a positive control for cell viability, cells grown in DMEM/F12 with 10 % SFB were also evaluated. Data are presented as means ± SEM of the percentage of ethidium incorporation (arbitrary units of fluorescence). Data from 3 cell lines and experiments were performed in duplicates (a), (b) and quadruplicates (c). *P < 0.05; **P < 0.01; ***P < 0.001; ****P < 0.0001; One-way ANOVA followed by Dunnett’s post hoc test. ns – Non-significant.

## 4. Discussion

In this study, human iPSC-derived astrocytes were characterized with regard to their responsiveness to short and long-term inflammatory conditions. Thus, human iPSC-derived astrocytes from healthy subjects were challenged with the pro-inflammatory cytokine TNF-α, which is able to initiate and sustain long-term features of astrogliosis (Cui, Huang, Tian, Zhao, & Zheng, 2011).

In glial cells, NF-kB remains in its physiologically latent cytoplasmic form, being translocated to the nucleus under inflammation (Kaltschmidt & Kaltschmidt, 2009). We noticed that human iPSC-derived astrocytes were capable of responding to short term TNF-α exposure through the translocation of NF-kB to the nucleus, a predicted event that occurs within an hour after inflammatory stimuli (Nelson et al., 2004; Romano, Freudenthal, Merlo, & Routtenberg, 2006).

Secretome analysis revealed that high levels of TNF-α were detected in the conditioned media 24 h after exposing cells to this cytokine, but it was difficult to distinguish between residual levels arising from the stimulus or secretion. However, TNF-α was almost undetectable in the conditioned medium and cell extracts from non-stimulated astrocytes, confirming that TNF-α is essentially produced by stimulated astrocytes. Since TNF-α via NF-kB activation triggers inflammatory events such as cytokines production, we also found that upon short-term TNF-α stimulation (24 h), either modulatory cytokines IL-2, IL-4 and IL-6 or the chemotactic and inflammatory cytokine IL-8 had their levels increased from pg/mL to ng/ml range, which is typical for inflammatory responses or pathological conditions (Stenken & Poschenrieder, 2015). However, IFN-γ and BDNF were also increased in the conditioned medium from astrocytes exposed to TNF-α, but to a lesser extent. It is important to take into account that over the course of acute inflammatory stimuli many factors and cytokines are expressed in a coordinated manner. Since IFN-γ is a pro-inflammatory factor, increases in its secretion may be important to induce and sustain astrogliosis (Yong et al., 1991). In a previous report, cultured rat astrocytes were exposed to different inflammatory stimuli, but only TNF-α was able to increase expression and secretion of BDNF via NF-kB activation (Saha, Liu, & Pahan, 2006). Even though the activity-dependent secretion BDNF is well documented in neuronal cells, the secretory nature of astrocytes suggests that increased BDNF secretion by TNF-α might be an attempt of stimulated-astrocytes to provide a neurotrophic factor that will rescue viability following acute inflammatory stimuli.

Interestingly, relatively high levels for IL-6 and IL-8 were detected in the conditioned medium from non-stimulated astrocytes (1.6 and 1.0 ng/ml, respectively). Of note, astrocytes are a major source of IL-6, which is important for synapse formation (Wei et al., 2011), maturation of dendritic spines (Wei et al., 2012) and sprouting of glutamatergic connections (Menezes et al., 2016), while IL-8 regulates angiogenesis by enhancing survival and proliferation of endothelial cells (Li, Dubey, Varney, Dave, & Singh, 2003).

As expected for mature astrocytes, almost all human iPSC-derived astrocytes were immunoreactive for GFAP and vimentin, though not all astrocytes necessarily express GFAP in the human brain tissue (Kettenmann & Verkhratsky, 2011; Kimelberg, 2004). Both GFAP and vimentin are considered markers of astrocytes and their overexpression characterizes primary features of reactive astrogliosis, since their deletion reduces reactive gliosis and promotes synaptic regeneration (Wilhelmsson et al., 2004). TNF-α is fairly known to induce overexpression of GFAP and vimentin (Perriot et al., 2018; Zhang et al., 2016) and our human iPSC-derived astrocytes were responsive to long-term TNF-α exposure by increasing GFAP and vimentin. The long-term exposure to TNF-α was also able to alter astrocyte morphology *in vitro*, since they displayed an elongated shape, namely polarized astrocytes, resembling those in response to a wound (Peng & Carbonetto, 2012) or recruited to mold the glial scar around injured sites (Adams & Gallo, 2018).

It is well documented the crucial role of astrocytes in maintaining glutamate homeostasis by continuous removal of extracellular glutamate from the synaptic cleft by glutamate transporters, a mechanism that avoids glutamate excitotoxicity (Rothstein et al., 1996). Recent evidences have shown that extracellular glutamate stimulates glutamate release from astrocytes in order to coordinate neuronal activity, pointing to an emerging role for astrocytes in modulating both glutamate uptake and release and extending their therapeutic potential (Mahmoud, Gharagozloo, Simard, & Gris, 2019). It could be noted that iPSC-derived astrocytes exposed to short term TNF-α had already showed impairment in the aspartate uptake. As astrogliosis progresses due to long term exposure to TNF-α, the aspartate uptake impairment worsened. Importantly, the impairment of glutamate uptake by prolonged exposure to TNF-α was not related to astrocyte viability, which confirms that they were dysfunctional indeed. Since atrophic astrocytes have been proposed to be dysfunctional, the impairment of glutamate uptake triggered by TNF-α intertwines both events in the context of astrocytopathy (Pekny & Pekna, 2016). Astrogliosis is considered a complex response, which varies from hypertrophy, atrophy, alteration in astrocytes biomarkers and dysfunction. Our findings show that human iPSC-derived astrocytes can be functionally activated, mimicking previously reported physiological aspects of astrogliosis described in animal models, human primary cultures and post-mortem brain studies. Astrogliopathology comprises both reactive astrogliosis and astrocytopathy that can co-exist after an insult by a series of changes that appear to be context- and disease-driven (Kim, Healey, Sepulveda-Orengo, & Reissner, 2018; Pekny & Pekna, 2016). These changes vary over time with a continuum of progressive alterations that can result in either beneficial or detrimental effects on the surrounding tissue (Burda & Sofroniew, 2014).

The inflammatory processes share similarities and differences between animal and human models, causing intense debate over the theme (Cauwels, Vandendriessche, & Brouckaert, 2013; Warren et al., 2015). For example, genomic responses to inflammatory challenges are divergent between both species, although the same dataset has been subject of contradictory interpretations in two different studies (Kodamullil et al., 2017; Mestas & Hughes, 2004; Seok et al., 2013; Takao & Miyakawa, 2015). Besides, signaling pathways of neuroinflammation share similarities between rodents and humans, but substantial differences were found for molecular and cellular interactions (Kodamullil et al., 2017). Astrogliosis has been systematically related to several CNS pathologies and the implementation of functional *in vitro* models of human neural cells has been subject of research in the last years. Therefore, a complete screening of inflammatory responses to different stressing stimuli by human astrocytes may shed light in the pathophysiology of neurodegenerative and psychiatric diseases. For instance, human iPSC-based model of astrogliosis may improve the understanding of neuroinflammation-related diseases and promote the development of new therapeutic strategies.

## Acknowledgements

We thank Fernanda Albuquerque, Gabriela Lopes Vitória, Ismael Carlos da Silva Gomes, Jarek Sochacki, Marcelo do Nascimento Costa and Renata Maciel Santos for technical support. The authors declare no competing financial interests. This work was sponsored by Fundação de Amparo a Pesquisa do Estado do Rio de Janeiro (FAPERJ), Conselho Nacional de Desenvolvimento Científico e Tecnológico (CNPq), Coordenação de Aperfeiçoamento de Pessoal de Nível Superior (CAPES), Instituto Nacional de Neurociência Translacional (INNT) and Banco Nacional de Desenvolvimento (BNDES), in addition to intramural grants from D’Or Institute for Research and Education.

---

Supplementary Figure 1 - Expression of housekeeping genes RPLP0, GAPDH and IPO8 was not affected by TNF-α exposure in human iPSC-derived astrocytes. Samples used in these experiments were also used in data presented on Figures 4b and 4c. Results were normalized by average values of all three housekeeping genes. Graphs represent data from 4 cell lines in experiments performed in triplicates. ns non-significant. Data are presented as means ± SEM.

## Author Contributions

P.T. improved the protocol for obtaining and culturing of human iPSC-derived astrocytes, performed cell morphology analyses and experiments for astrogliosis characterization. P. T. and E. C. L. cultured and adjusted human iPSC-derived astrocytes for all performed experiments. P.T., J. A. S. and I. M. O. designed and analyzed NF-kB experiments. J. G., C. T. R, J. C. F. M and D. P. G., performed and analyzed secretome data. P. L. C., S. D. and F. M. R. performed and analyzed quantitative real time PCR experiments. P.T., E.C.L. and A. L. M. V. conducted D-aspartate uptake assay and analysis. P. F. L. and P. L. C. designed schematic figures. P.T. and S. K. R. designed the research. P.T., P.F.L, J.A.S., L.O.P. and S.K.R. analyzed data, discussed results and wrote the paper. All authors discussed results and validation steps.

