## Supplementary figures and images for "Short and long TNF-alpha exposure recapitulates canonical astrogliosis events in human induced pluripotent stem cells-derived astrocytes"

### Supplementary Figure 1

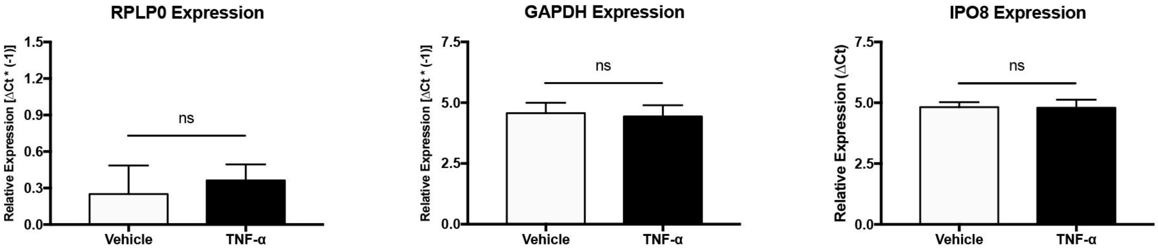
